# Elicitation of superinfection exclusion by p28 of turnip crinkle virus is separable from its replication function with mutations at specific amino acid residues

**DOI:** 10.1101/596767

**Authors:** Qin Guo, Shaoyan Zhang, Rong Sun, Xiaolong Yao, Xiao-Feng Zhang, Satyanarayana Tatineni, Tea Meulia, Feng Qu

**Affiliations:** Department of Plant Pathology, The Ohio State University. Wooster, OH 44691; Fujian Key Laboratory of Plant Virology, Institute of Plant Virology, Fujian Agriculture and Forestry University, Fuzhou, China; USDA-ARS and Department of Plant Pathology, University of Nebraska-Lincoln, Lincoln, NE 68583; Molecular and Cellular Imaging Center, Ohio Agricultural Research and Development Center, The Ohio State University. Wooster, OH 44691

**Author notes:** Primary contact: Feng Qu.

## Abstract

We recently reported that the p28 auxiliary replication protein encoded by turnip crinkle virus (TCV) is also responsible for eliciting superinfection exclusion (SIE) against superinfecting TCV. However, it remains unresolved whether the replication function of p28 could be separated from its ability to elicit SIE. Here we report the identification of two single amino acid (aa) mutations that decouple these two functions. Using an *Agrobacterium* infiltration-based delivery system, we transiently expressed a series of p28 deletion and point mutants, and tested their ability to elicit SIE against a co-introduced TCV replicon. We found that substituting alanine (A) for valine (V) and phenylalanine (F) at p28 positions 181 and 182, respectively, modestly compromised SIE in transiently expressed p28 derivatives. Upon incorporation into TCV replicons, V181A and F182A decoupled TCV replication and SIE diametrically. While V181A impaired SIE without detectably compromising replication, F182A abolished TCV replication but had no effect on SIE once the replication of the defective replicon was restored through complementation. Both mutations diminished accumulation of p28 protein, suggesting that p28 must reach a concentration threshold in order to elicit a strong SIE. Importantly, the severe reduction of F182A protein levels correlated with a dramatic loss in the number of intracellular p28 foci formed by p28-p28 interactions. Together these findings not only decouples the replication and SIE functions of p28, but also unveils a concentration dependence for p28 coalescence and SIE elicitation. These data further highlight the role of p28 multimerization in driving the exclusion of secondary TCV infections.

**IMPORTANCE:** Superinfection exclusion (SIE) insulates virus-infected cells from subsequent invasion by the same or closely related viruses. SIE has been observed in both animal and plant virus-infected cells. Therefore, a thorough understanding of how SIE is achieved at the molecular level is expected to inspire novel strategies for combating virus infections in humans, animals, and plants. Our group has been using turnip crinkle virus (TCV) to elucidate the molecular interactions critical for SIE elicitation. The current study builds on the previous observation that TCV SIE is elicited by one single TCV-encoded protein (p28), and further identifies key regions and amino acids that are needed for SIE. We unravel key amino acid changes that decouple the replication and SIE functions of p28, and provides novel mechanistic insights of SIE.

## INTRODUCTION

Many viruses block the subsequent entry and/or replication of the same or closely related viruses in the cells they occupy, through a process known as superinfection exclusion (SIE). These viruses include important human and animal pathogens such as human immunodeficiency virus (HIV), hepatitis C virus (HCV), West Nile virus (WNV), and vesicular stomatitis virus (VSV) (1–5), as well as plant pathogens like tobacco mosaic virus (TMV), citrus tristeza virus (CTV), cucumber mosaic virus (CMV), wheat streak mosaic virus (WSMV), turnip crinkle virus (TCV) (6–11), and more recently sonchus yellow net virus (12). SIE in plant virus infections is also thought to underlie the well-documented cross protection phenomenon (13–15). Cross protection insulates plants from a disease-causing virus by simply pre-inoculating them with a mild variant of the same virus. Although cross protection has been adopted for plant virus disease management for at least 50 years (13), its underlying molecular mechanism remains poorly understood.

A number of earlier studies investigated SIE of plant viruses by tagging the same virus with two different fluorescent proteins (e.g. green fluorescent protein [GFP] and mCherry) (6, 7, 9, 10, 16, 17). These studies demonstrated that most of the viruses excluded their own variants at the single cell level, thus implicating intracellular exclusion as one of the primary causes for SIE. Using this approach, Zhang and colleagues (16) found that SIE in TCV infections occurred inside individual cells, and it acted to block the replication of the superinfecting TCV replicons. Strikingly, this replicational blockage was mediated by p28, one of the TCV-encoded proteins required for TCV genome replication (16). The fact that p28, the auxiliary replication protein of TCV, functions to both facilitate and repress the replication of TCV genomic RNA (gRNA), along with the observation that fluorescent protein-tagged p28 formed large intracellular bodies that coalesced other p28 variants, prompted the hypothesis that p28 undergoes concentration-dependent conformational changes to accommodate its dual functions. Namely, the p28 protein assumes one conformation at lower concentrations to enable replication, but another self-perpetuating conformation at higher concentrations to block replication (16, 18).

TCV is a small icosahedral plant virus with a (+)-strand RNA genome that encodes five proteins. The 5’ proximal p28 and its C-terminally extended derivative – p88 – are both translated directly from TCV genomic RNA (gRNA), and are themselves required for gRNA replication (Fig. 1A). Since the production of p88 depends on infrequent translational read-through of the p28 stop codon (UAG), the intracellular concentration of p28 far exceeds that of p88 (19). However, p28 lacks the conserved RNA-dependent RNA polymerase (RdRP) motif, is hence known as the auxiliary replication protein. On the other hand, p88 is thought to be directly responsible for gRNA replication through its RdRP activity. TCV additionally encodes two small movement proteins, p8 and p9, that are translated from subgenomic RNA 1 (sgRNA1), and a capsid protein/silencing suppressor from sgRNA2 (Fig. 1A) (20–22).

**Figure 1.**
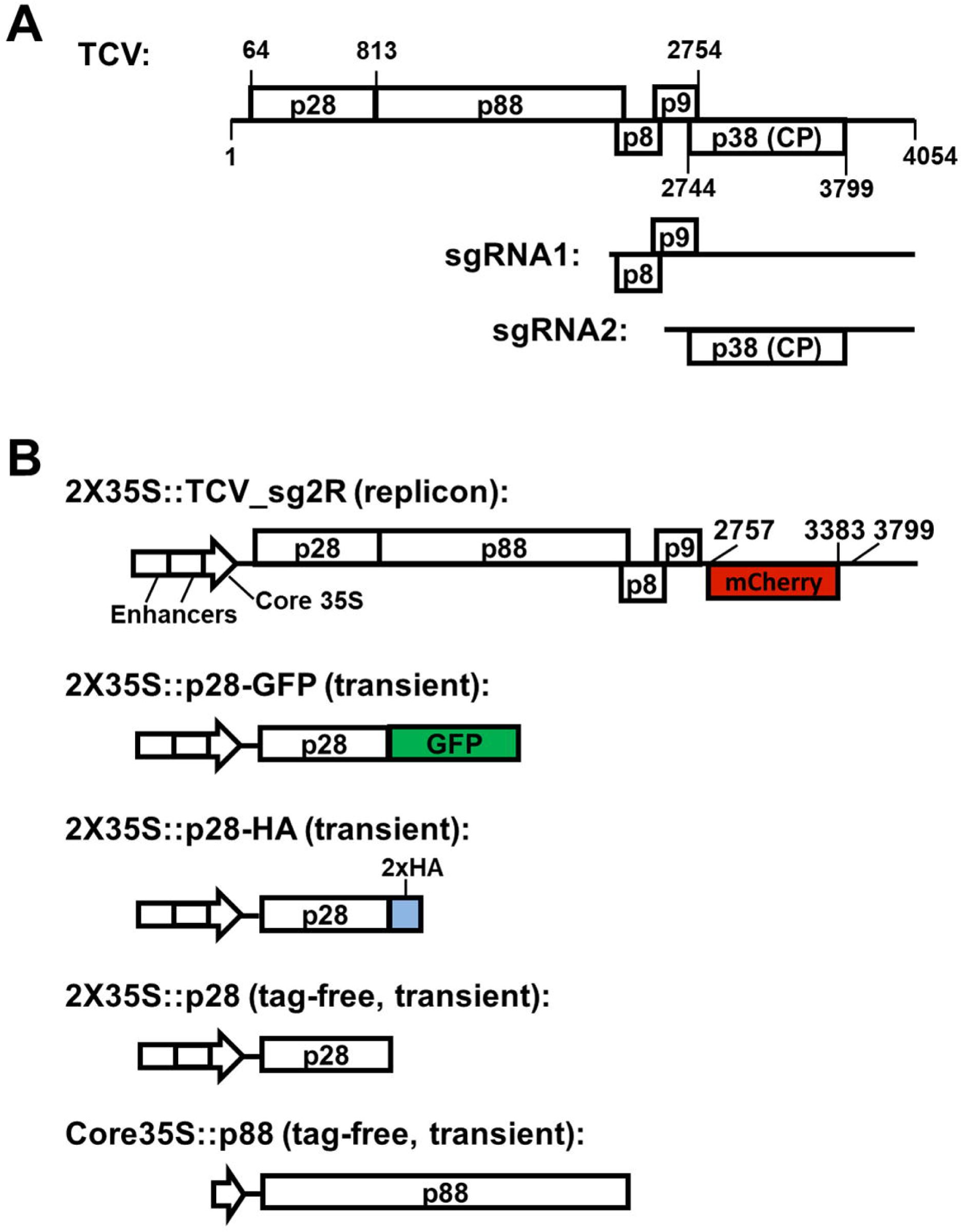
TCV genome organization, and the configurations of the constructs used in this study**. A:** the organization of the 4,054 nt single-stranded, positive sense (+) TCV genome. The various boxes with labels within (e.g. p28, p88, p8, p9, p38) denote the proteins encoded by the TCV genome, with the boundaries of some ORFs shown. The three 3’ proximal proteins (p8, p9, and p38) are translated from two subgenomic RNAs (sgRNA1 and 2). **B**: representative constructs used in this study. Most constructs were equipped with the strong 35S promoter with the enhancer duplicated (2X35S) in order to ensure efficient transcription. Only the last, p88-expressing construct was driven by the relatively weak core 35S promoter as high level of p88 translation was shown to repress TCV replication (23). The top most construct is a replicon shown previously to launch TCV replication in plant cells, leading to mCherry expression from the sgRNA2 (20, 36). All other constructs direct transient expression of the respective proteins in the absence of TCV replication.

In the recent study by Zhang and colleagues (16), we established that p28 was necessary and sufficient to elicit SIE against a co-introduced TCV replicon, and that it likely accomplished this by sequestering freshly translated p28 molecules into self-perpetuating p28 polymers. These earlier observations raised at least three questions: (i) does the SIE activity of p28 require any specific domains or amino acid residues in p28? (ii) can this replication-repressing SIE activity of p28 be decoupled from its essential role in replication? (iii) can either of the p28 functions (replication and SIE) be decoupled from the RdRP activity of p88? This last question is of interest as p88 encompasses the entire p28 at its N-terminal 2/5 (Fig. 1A). The current study addresses these questions by identifying a 12 amino acid (aa) region in p28 that is required for efficient SIE. Moreover, two single aa changes within this region not only enabled the decoupling of SIE activity of p28 from its replication function, but also the separation of the replication function of p28 from that of p88. Finally, both single aa mutations lowered the cellular concentration of p28, and diminished the number of p28 inclusion bodies. Our findings unveil novel mechanistic details of SIE elicitation by TCV p28, and provide powerful new tools for further elucidation of TCV SIE.

## RESULTS

### Deletion mutagenesis of p28-GFP fusion protein identifies a small region needed for robust SIE elicitation

We recently reported that a C-terminal GFP fusion bolsters the ability of p28 to elicit SIE to TCV (16). Specifically, expression of the p28-GFP fusion protein in *Nicotiana benthamiana* cells abolished the replication of TCV_sg2R, an mCherry-expressing TCV replicon, in the same cells, erasing the replication-dependent mCherry fluorescence (16). Additionally, the strong replicational repression by p28-GFP correlated with the occurrence of large, intense green fluorescent foci in these cells (16). These easy-to-monitor fluorescent phenotypes prompted us to use p28-GFP as the launch pad to map domains in p28 essential for SIE elicitation. To systematically screen for these domains, we generated a series of in-frame deletions within the p28 portion of p28-GFP (Fig. 2A). The effect of these deletions on the SIE-eliciting ability of p28 was evaluated by expressing the mutant proteins in *N. benthamiana* cells, and assessing the replication of the co-introduced TCV_sg2R replicon with Northern blot hybridizations. In addition, we also monitored the replication-dependent diffuse mCherry fluorescence, as well as the GFP-fluorescent mutant fusion proteins, using confocal microscopy.

**Figure 2.**
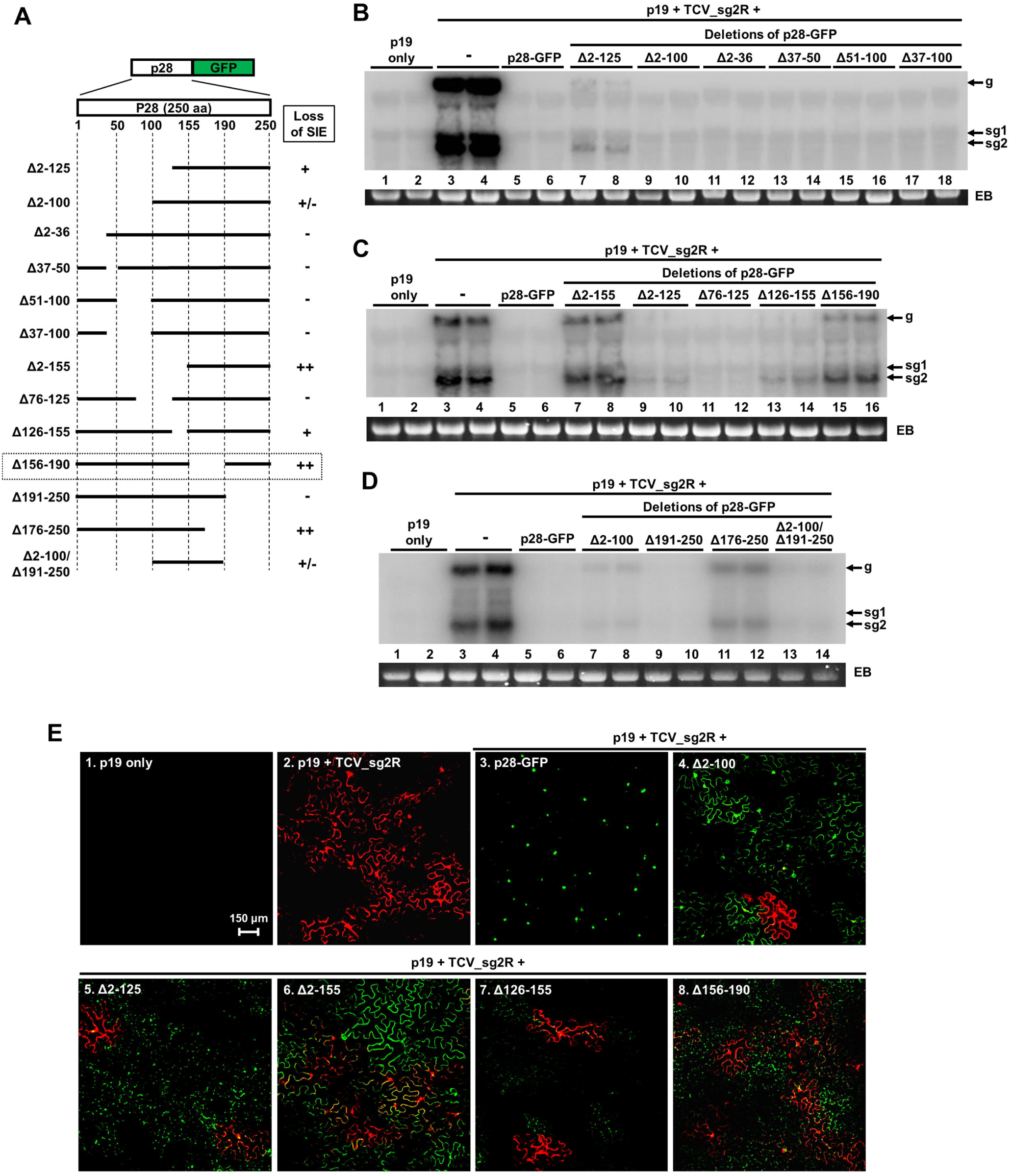
Deletion mutagenesis on the p28-GFP backbone identifies the aa 156-190 region as needed for robust SIE. **A:** diagrams of deletion mutants of p28-GFP showing their respective deleted regions. **B, C, D:** Northern blotting results showing the levels of replicational repression imposed on the TCV_sg2R replicon by p28-GFP and its deletion mutants. The p19 silencing suppressor was included to mitigate the impact of RNA silencing. The double loadings represented two independent samples extracted from different leaves. The positions of gRNA, sgRNA1 and 2 were highlighted with arrows (g, sg1, sg2). EB: ethidium bromide-stained gel. **E:** confocal images of *N. benthamiana* leaf cells treated with select p28-GFP deletion mutants along with the TCV_sg2R replicon.

As shown in Fig. 2B, deletion of the N-terminal half of the 250-aa p28 (mutant Δ2-125) caused only minimal loss in the SIE elicitation by p28-GFP, leading to a very modest recovery of TCV_sg2R replication (Fig. 2A; and Fig. 2B, lanes 7 and 8). A specific N-terminal domain could not be delineated because none of the smaller deletions within this region, including Δ2-36, Δ37-50, Δ37-100, Δ51-100, and Δ76-125, led to detectable replication of TCV_sg2R (Fig. 2B, lanes 11-18; Fig. 2C, lanes 11-12). These results suggest that, in the presence of the C-terminal GFP tag, the N-terminal half of p28 plays a modest role in SIE elicitation. Consistent with this conclusion, the Δ2-100 deletion had an even smaller effect than Δ2-125 on SIE, leading to only occasional detection of TCV_sg2R replication (Fig. 2B, lanes 9 and 10; Fig. 2D, lanes 7 and 8).

Interestingly, the Δ2-155 mutant, which extends the Δ2-125 deletion for just 30 aa, permitted a substantially higher level of TCV_sg2R replication (Fig. 2C, compare lanes 7-8 with 9-10). However, a separate deletion of the 30 aa alone (Δ126-155) had a much weaker, though consistently detected release of TCV_sg2R replication (Fig. 2C, compare lanes 13, 14 with lanes 7, 8). This suggested that loss of SIE activity by the mutant Δ2-155 could be attributed to the additive or synergistic effect of two regions – aa 2-125 and 126-155. Nevertheless, specific SIE-eliciting sequence motifs or aa residues within the first 155 aa region could not be reliably identified with the GFP-tagged constructs.

Surprisingly, another relatively small deletion (Δ156-190, 35 aa in total) caused a loss of SIE activity that rivaled the large Δ2-155 deletion (Fig. 2C, compare lanes 15, 16 with 7 and 8), thus identifying a region with a relatively strong effect. Beyond aa 190, the entire C-terminal 60-aa region could be removed without detectably compromising the SIE-eliciting activity of p28-GFP (Fig. 2D, Δ191-250, lanes 9 and 10). Indeed, a double deletion mutant combining the Δ2-100 and Δ191-250 deletions was no worse than Δ2-100 alone at repressing TCV_sg2R replication (Fig. 2D, compare lanes 13, 14 with lanes 7 and 8). This further confirmed that the last 60-aa region was dispensable for SIE elicitation by p28-GFP.

Notably, the loss of SIE activity by many of the deletion mutants was accompanied by dramatic changes in the intracellular behavior of mutant proteins. For example, although the SIE loss caused by the Δ2-100 deletion was minimal, the Δ2-100-GFP formed large foci in substantially fewer cells than the wild-type p28-GFP (Fig. 2E, panels 3 and 4), and instead exhibited a distribution resembling soluble GFP in many cells (Fig. 2E, panel 4). Nevertheless, the smaller, more numerous foci formed by the GFP-tagged Δ2-125, Δ126-155, and Δ156-190 proteins appear to be consistent with a requirement for larger aggregates in SIE elicitation (Fig. 2E, panels 5, 7, and 8). Further supporting this idea is the observation that the Δ2-155-GFP mutant protein was predominantly diffusely distributed, suggestive of a soluble protein (Fig. 2E, panel 6).

### Additional deletions within the aa 156-190 region identify a 12-aa stretch with a significant role in eliciting SIE

That a substantial loss of SIE-eliciting activity could be inflicted by a mere 35-aa deletion (Δ156-190) prompted us to further interrogate this small region using the p28-GFP fusion construct. To this end, three smaller deletions (Δ156-170, Δ171-182, Δ183-190, deleting 15, 12, and 8 aa, respectively) were generated and the resulting deletion mutants were analyzed for their ability to elicit SIE against TCV_sg2R. As shown in Fig. 3A, neither Δ156-170 (lanes 9 and 10) nor Δ183-190 (lanes 13 and 14) caused detectable loss of the SIE-eliciting activities. By contrast, deletion of aa 171-182 appeared to account for most of the SIE loss observed with the Δ156-190 mutant.

**Figure 3.**
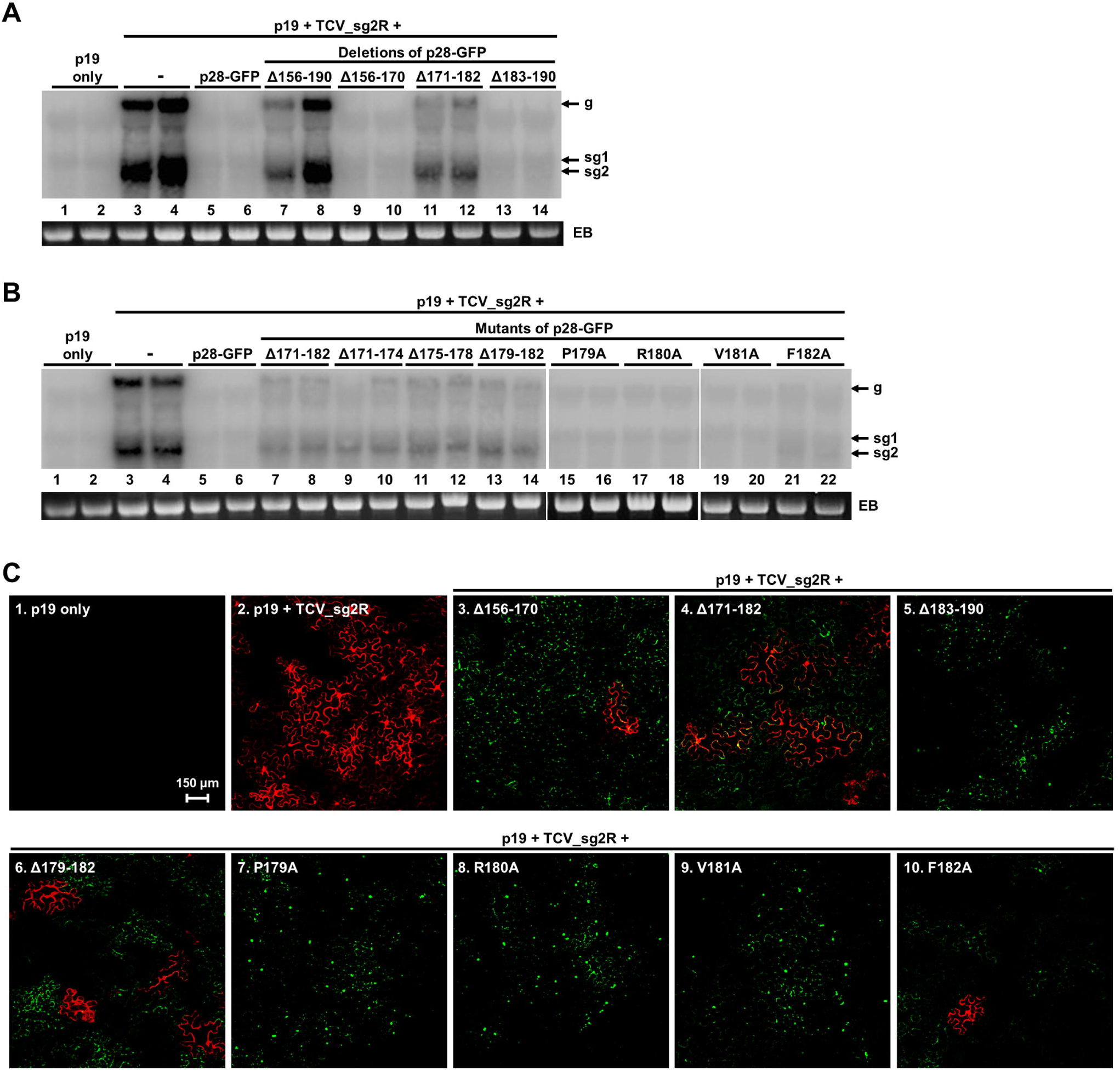
Mapping the aa 156-190 region of p28-GFP with smaller deletions and single aa exchanges identifies aa 171-182 as the key determinant of SIE. **A and B:** Northern blots showing the levels of replicational repression imposed on the TCV_sg2R replicon by various smaller deletion and single aa mutants of p28-GFP. **C**: confocal images of *N. benthamiana* leaf cells treated with select p28-GFP mutants along with the TCV_sg2R replicon.

We next investigated whether this 12-aa region contained any specific aa residue(s) critical for SIE elicitation by generating three consecutive 4-aa deletions. As shown in Fig. 3B, lanes 9-14, all three deletion mutants compromised the SIE-eliciting activity of p28-GFP to similar extents as the 12-aa deletion mutant Δ171-182 (lanes 7 and 8). Therefore, it appears that the entire 12-aa region is needed for eliciting a robust SIE. We then chose the last four aa residues of this region for single aa mutagenesis and changed each of them to alanine (Fig. 3B, lanes 15-22). When these single aa mutants were introduced into plant cells along with the TCV_sg2R replicon, only the F182A mutant caused a barely detectable accumulation of TCV_sg2R RNAs (Fig. 3B, lanes 21 and 22), suggesting that the phenylalanine to alanine change weakly compromised the SIE-eliciting ability of p28-GFP.

Confocal microscopy showed that Δ156-170, Δ171-182, and Δ183-190 all abolished the ability to form large aggregates (Fig. 3C, panels 3 and 4, also see Fig. 2E), and instead formed smaller, but more numerous foci that sometimes arranged along cell boundaries. The Δ156-170 mutant was noteworthy because it did allow occasional escape of red fluorescent cells, thus TCV_sg2R replication, even though such low-level replication was not detected with Northern blotting (see above). This suggested that it was important to use both Northern blotting and confocal microscopy to capture the full spectrum of SIE-eliciting activities by these mutants.

Interestingly, the smaller, 4-aa deletion mutant Δ179-182 and the larger, 12-aa deletion mutant Δ171-182 behaved highly similarly when inspected with confocal microscopy. Both permitted TCV_sg2R replication in easily detectable numbers of cells, even though their replication levels as measured with Northern blotting was much lower than controls without any p28-GFP derivatives (Fig. 2B, lanes 7-14). Finally, three of the four single aa mutants, P179A, R180A, and V181A, formed large aggregates very similar to those of wild-type p28-GFP, though smaller foci were also seen (Fig. 3C, panels 7-9). By contrast, in cells treated with the F182A mutant, large aggregates were absent (Fig. 3C, panel 10). Among these cells, replication-dependent red fluorescence was occasionally observed, thus confirming the weak TCV_sg2R replication detected with Northern blotting (Fig. 2B, lanes 21-22).

### Replacing the C-terminal GFP with a double HA tag unveils additional regions important for SIE-elicitation by p28

Experiments using the p28-GFP fusion protein mutants identified the 12-aa region spanning aa 171-182 as critically important for p28 to elicit SIE. But they failed to identify any other p28 regions that were specifically needed for SIE elicitation. These results contrasted with our previous observations implicating the N-terminus of p28 in SIE elicitation. We showed earlier that fusing a short G11 tag (25 aa) at the N-terminus rendered p28 incapable of eliciting SIE (16). We also found that a truncated form of p28 missing the first 36 aa was no longer able to elicit SIE (16, 23). Did the C-terminal GFP fusion in p28-GFP allow p28 to circumvent certain other critical regions, possibly through the tendency of GFP to self-dimerize (24)? To address this question, we decided to adopt p28-HA, another strong SIE-eliciting p28 variant with a C-terminal, duplicated hemagglutinin tag (16, 23). To this end, the GFP in all deletion mutants described earlier was replaced with the HA tag and the resulting HA-tagged mutants were retested for SIE induction.

As shown in Fig. 4A, the HA substitution indeed revealed a role for the N-terminal region in SIE elicitation. The SIE-eliciting activity of p28-HA was not detectably weakened by deleting aa 51-100 or 191-250, suggesting that up to 110 amino acid residues of p28 could be removed without detectably weakening SIE elicitation. However, this activity is clearly compromised by deletions of aa 2-36, 126-155, and 156-190, and aa 37-100, 76-125, and 183-190 to lesser extents. Therefore, use of p28-HA as the backbone permitted the identification of two additional p28 regions with potentially specific roles in SIE elicitation, namely aa 2-50, and 126-155.

**Figure 4.**
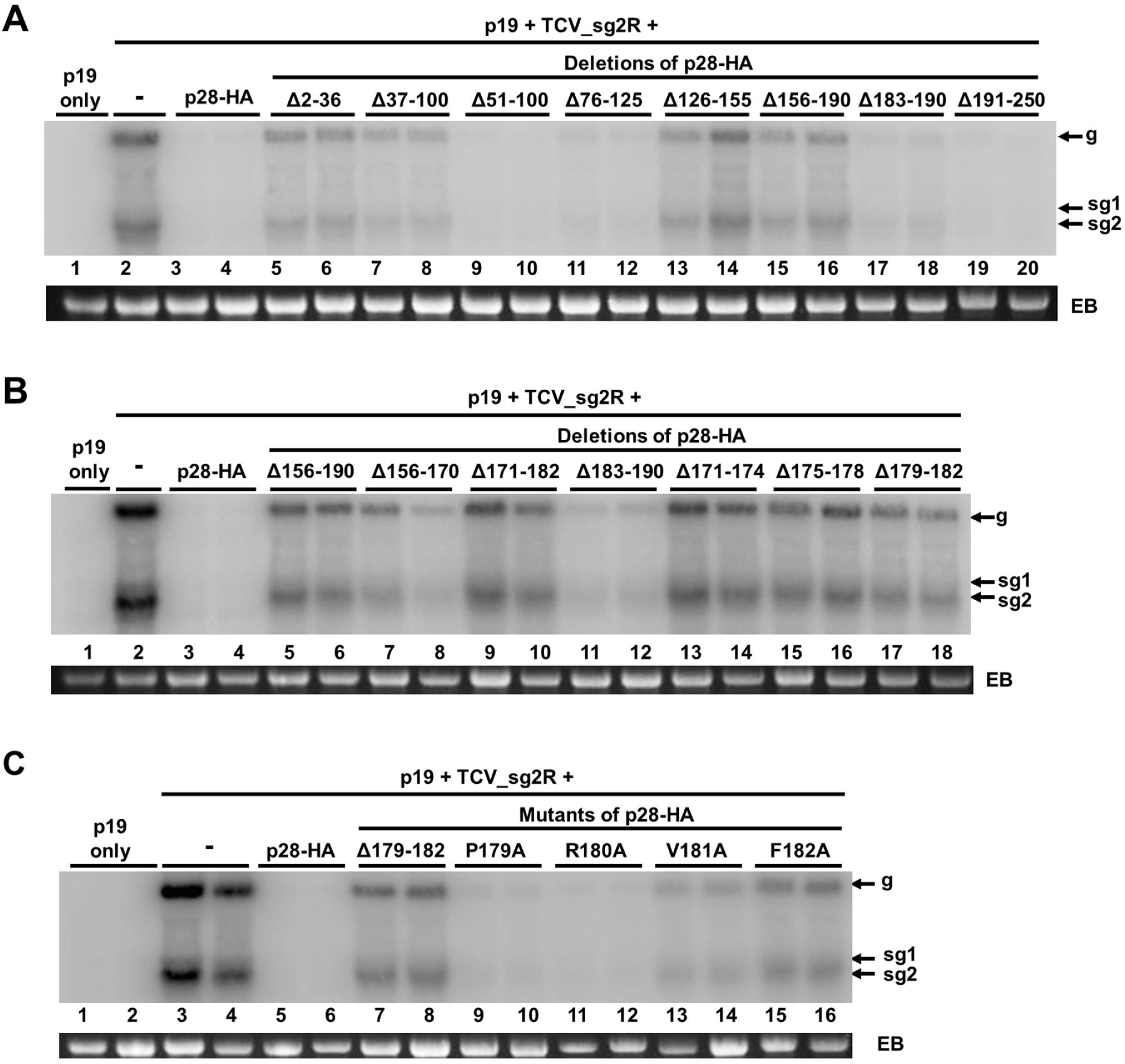
Use of the p28-HA backbone reveals additional regions contributing to SIE elicitation by p28. All panels are Northern blots showing the levels of replicational repression imposed on the TCV_sg2R replicon by various mutants of p28-HA.

### Use of p28-HA reveals key amino acid residues within the aa 171-182 region critical for SIE elicitation

We next attempted to fine-map the 171-182 region, whose deletion compromised SIE in both GFP and HA-tagged p28 variants, using HA-tagged mutants. As shown in Fig. 4B, use of the p28-HA backbone revealed partial SIE losses caused by the Δ156-170 and Δ183-190 deletions (lanes 7, 8, 11, 12) that were undetectable with the p28-GFP backbone (Fig. 3A, lanes 9, 10, and 13, 14). However, it is important to note that only the deletion of the aa 171-182 core sequence fully recapitulated the phenotype of the larger 156-190 deletion (Fig. 4B, compare lanes 5, 6 with 9, 10). Furthermore, each of the three smaller, 4-aa deletions (Δ171-174, Δ175-178, and Δ179-182) within the aa 171-182 region inflicted an SIE loss that is nearly identical to the 12-aa Δ171-182 (Fig. 4B, lanes 13-18).

We next re-examined the effect of four single aa substitutions using the p28-HA backbone. As shown in Fig. 4C, the very weak SIE loss caused by the F182A mutation in p28-GFP background now became easily detectable (lanes 15 and 16). Indeed, only the R180A mutation failed to inflict a meaningful loss in the SIE-eliciting activity of p28-HA (lanes 11 and 12). By contrast, both P179A and V181A mutations correlated with weak losses in SIE elicitation, with P179A being slightly less disruptive than V181A (lanes 9, 10, 13 and 14).

### TCV replicons containing the V181A mutation overcome SIE in a substantial fraction of cells

We next wondered whether the single-aa mutations that compromised SIE in transiently expressed p28-GFP and p28-HA had similar effects in TCV replicons. To resolve this question, we introduced the V181A and F182A mutations into TCV_sG6H and TCV_sg2R, two TCV replicons that encode GFP [sG6H denotes GFP with an N-terminal 6XHis tag (25)] and mCherry in place of CP, respectively (Fig. 1). These replicons were then tested in co-infected cells to determine whether the mutations compromised SIE of TCV.

Although the GFP-encoding replicon (TCV_sG6H) appeared to replicate less robustly, and produced much less sgRNAs (Fig. 5A, lane 2), it produced easily detectable green fluorescence (Fig. 5B, top left), probably due to translational initiation from an internal ribosomal entry site identified recently (26). Replicons with the V181A mutation (V181A_sG6H and V181A_sg2R) replicated to levels comparable to wild-type replicons encoding wild-type p28 (and p88) (Fig. 5A, compare lanes 4, 5 with 2, 3, respectively). By contrast, the F182A mutation abrogated TCV replication to levels below the detection limit of Northern blotting (lanes 6 and 7). Next, the V181A replicons were subjected to co-infection experiments to assess the levels of SIE. As shown in Fig. 5B, while wild-type replicons exhibited strong SIE that precluded the co-expression of GFP and mCherry in the same cells (top row), the V181A replicons permitted such co-expression in approximately 19% of green- or red-fluorescent cells. These results demonstrated that the same V181A mutation that compromised the ability of p28-HA to repress TCV replication also compromised the ability of p28 to elicit SIE in TCV replicons.

**Figure 5.**
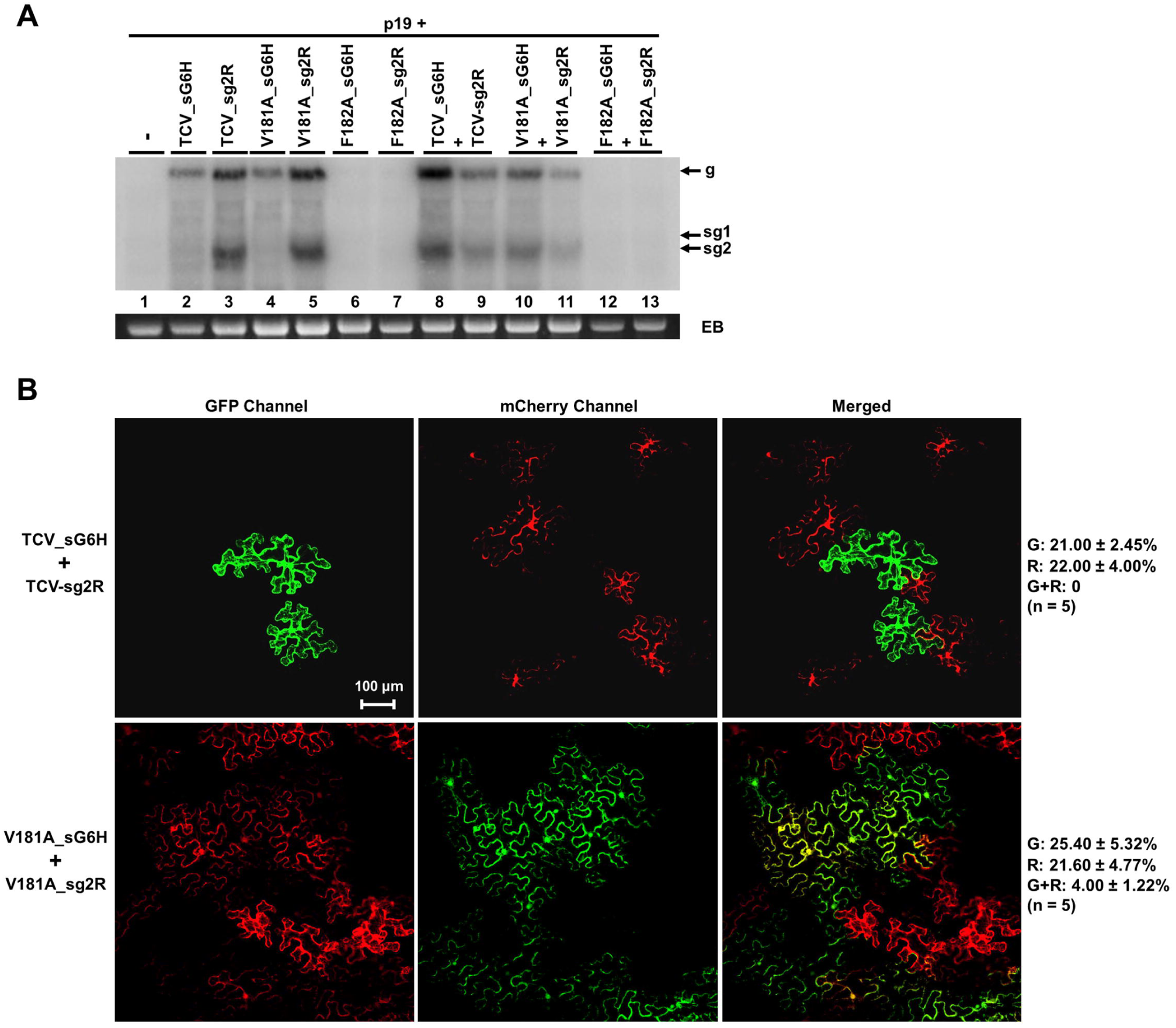
Replication and SIE of mutant replicons with the V181A and F182A mutations. **A**: Northern blot showing the replication (lanes 4-7) and SIE (lanes 10-13) of the mutant replicons encoding different fluorescent proteins (GFP or mCherry). **B**: confocal images showing robust SIE between two wild-type replicons (top panels), and the partially compromised SIE between the two V181A mutant replicons. The quantification was based on five independent leaf areas each containing 40-50 cells. See Materials and Methods for details. G = GFP-fluorescent cells, R = mCherry-fluorescent cells, G+R = cells containing both GFP and mCherry.

### The F182A mutation disrupts the replication function of p28 but not p88

The impact of the F182A mutation on replicon SIE could not be readily examined because it abolished TCV replication when incorporated in replicons (Fig. 5). Since the p28 coding sequence is shared by p88 in the replicon context, the F182A change could have disrupted the replication function of p28, or p88, or both. To resolve this issue, we attempted to complement the replication of the F182A_sg2R replicon using transiently expressed, tag-free p28 and p88 (Fig. 1B). As shown in Fig. 6A, F182A_sg2R replication was restored to easily detectable levels by transiently expressed p28 (compare lanes 9, 10 to 5, 6), but not p88 (lanes 13 and 14). Therefore, the F182A mutation primarily disrupted the auxiliary replication function of p28. While minor perturbation of p88 function by F182A could not be ruled out, such perturbation did not abolish TCV replication. Taken together, the F182A mutation severely debilitated the replication function of p28, but not p88. In conclusion, this mutation decouples the replication functions of p28 and p88.

**Figure 6.**
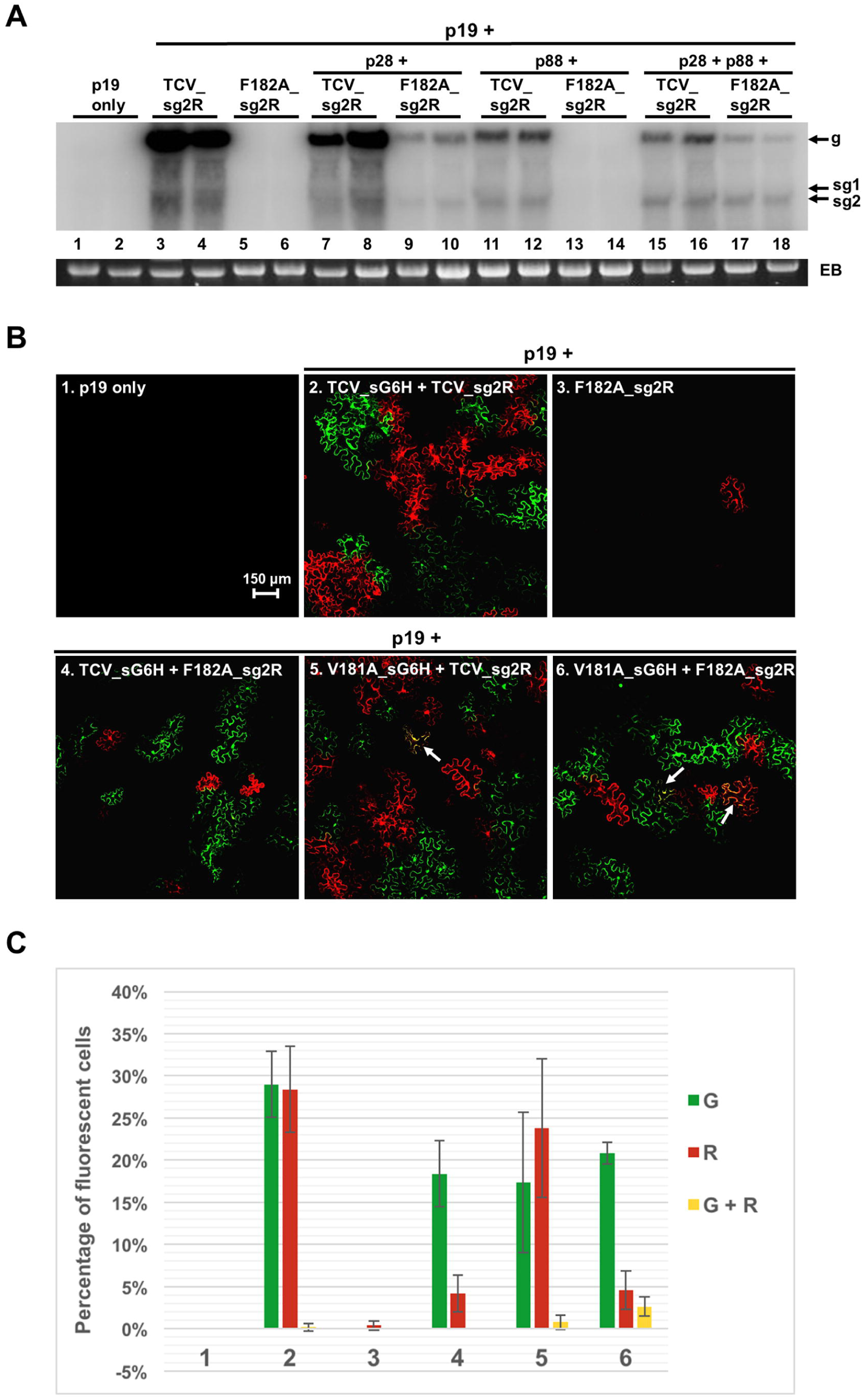
In replicons, the F182A mutation selectively abrogates the replication function of p28, but not its ability to elicit SIE. **A:** Northern blot showing that the replicational defect of the F182A replicon was complemented by transiently expressed p28 but not p88. **B:** confocal images showing that the replicational defect of the F182A replicon was also complemented by co-introduced wild-type and V181A replicons, and the successfully complemented F182A replicon exerted robust SIE against the wild-type replicon. **C**: quantification of replicational complementation and SIE of the F182A replicon. The numbering of the bar groups corresponds to that of Fig. 6B panels. In all sets leaf areas of the same size, containing 40-50 cells each, were chosen from five different leaves and counted for cells with different fluorescence (GFP, mCherry, or both).

### An F182A-containing replicon, upon replicational complementation, exerts robust SIE against a wild-type replicon

We next examined whether the replicational defect of the F182A-containing replicons could be complemented by another replicon encoding a replication-competent p28, and whether the p28(F182A) protein produced by the complemented replicon might exert SIE against the replicon that provided the complementing p28. To answer this question, we paired the F182A_sg2R replicon with TCV_sG6H or V181A_sG6H, in co-infections of *N. benthamiana* cells. As shown in Fig. 6B, panels 4 and 6, both TCV_sG6H and V181A_sG6H replicons complemented the F182A defect, enabling an approximately ten-fold increase in the number of cells replicating F182A_sg2R (Fig. 6B, panels 3, 4, and 6; Fig. 6C, red bars in groups 3, 4, and 6).

However, complementation by the two GFP-encoding replicons led to starkly different SIE phenotypes. The complementation by TCV_sG6H, which encodes a wild-type p28, gave rise to mCherry-expressing cells that never contained GFP fluorescence (Fig. 6B, panel 4; Fig. 6C, group 4). Conversely, the GFP-expressing cells also never contained mCherry. Because the TCV_sG6H construct must reside in the same cells as the F182A_sg2R construct in order to supply the replication-complementing wild-type p28, these observations strongly suggested that the p28(F182A) protein, now produced by the replicating F182A_sg2R replicon, exerted a powerful SIE to block TCV_sG6H replication. Note that we have shown in a previous study (16) that commencement of TCV replication in different agro-infiltrated *N. benthamiana* cells was highly asynchronous, meaning many cells could translate wild-type p28 protein from the 2X35S-driven replicon transcripts, but fail to immediately embark on replicon replication. These cells would provide the p28 protein to complement the replication of F182A_sg2R, the latter then turn around to exert SIE against the p28-providing wild-type replicon.

The assertion that both F182A_sg2R and TCV_sG6H constructs must have been present in those red fluorescent cells is further bolstered by the contrasting SIE phenotype observed from the complementation mediated by V181A_sG6H (Fig. 6B, panel 6; Fig. 6C), as these experiments were carried out in parallel. Specifically, when the replication of F182A_sg2R was complemented with the V181A_sG6H replicon, approximately one half of the red fluorescent cells also contained GFP fluorescence. This probably reflected the partial loss of SIE-eliciting capacity by the p28(V181A) protein (Fig. 5B. Also see Fig. 6B, panel 5). But this result also demonstrated that two different replicon constructs readily entered the same agro-infiltrated cells. Together these results indicated that when expressed from a replicating replicon, the p28(F182A) protein is fully capable of exerting a strong SIE against another replicon in the same cells, despite its inability to facilitate replication on its own. Therefore, in addition to decoupling the replication function of p28 and p88, the F182A mutation also decoupled the SIE function and replication function of p28 – but in a manner diametrical to the V181A mutation.

### The p28(V181A) and p28(F182A) mutant proteins accumulated to substantially lower levels in cells, and formed fewer p28 foci

Why did the V181A mutation weaken SIE in both transiently expressed p28 proteins and TCV replicons, whereas F182A did so only in transiently expressed p28? To address this question, we assessed the accumulation levels of the mutant p28 proteins using Western blotting. We chose to assay the GFP and HA-tagged forms of p28 because a p28-specific antibody is not yet available despite repeated attempts using multiple vendors. As shown in Fig. 7A and B, both GFP and HA-tagged forms of V181A and F182A mutant p28 accumulated to levels substantially lower that their wildtype p28 counterparts. Importantly, while the HA-tagged forms of both mutant proteins diminished to levels below the detection limit (Fig. 7B), the GFP-tagged forms showed a slight difference between the two, with V181A, but not F182A, being marginally detectable. These results suggested that both mutations compromised p28 accumulation, but the effect of F182A was more pronounced.

**Figure 7.**
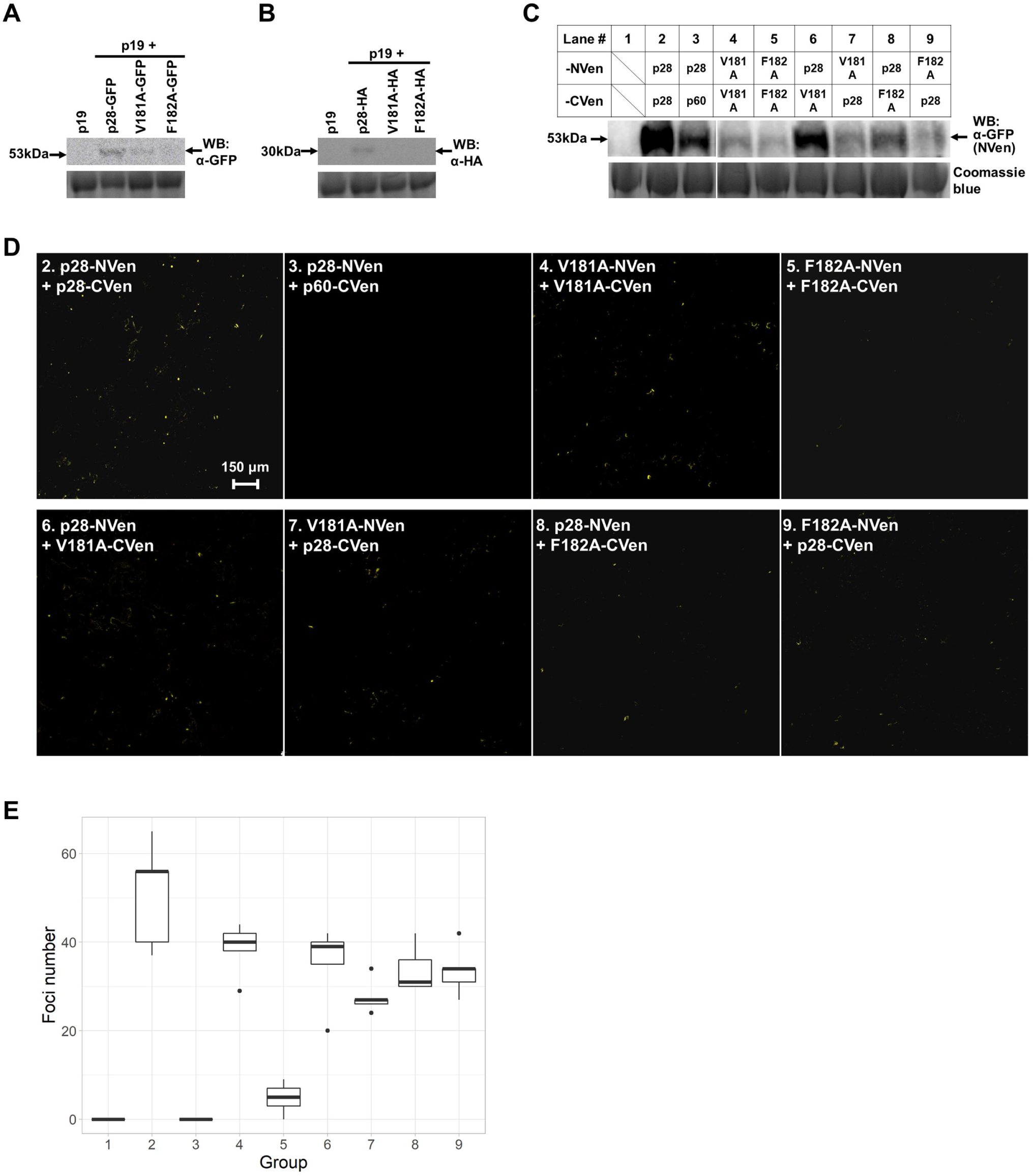
The V181A and F182A mutations reduce the level of transiently expressed p28 proteins, and the number of polymeric foci formed by interacting p28 proteins. **A**: Western blot showing accumulation levels of GFP-tagged wildtype p28 protein and its V181A and F182A mutants. **B**: Western blot showing accumulation levels of HA-tagged wildtype p28 protein and its V181A and F182A mutants. **C**: Western blot showing the accumulation levels of NVen-tagged p28, p28)V181A), p28(F182A), in the presence of various CVen-tagged interaction partners. **D**. Confocal images of multimeric p28 foci formed through p28-p28 interactions, visualized using mVenus-based BIFC. The numbering of the panels corresponds to that of lanes in Fig. 7C. **E.** Quantification of the number of p28 foci observed in various BIFC combinations. See Materials and Methods for details of quantification methods. The numbering of the groups corresponds to that of lanes in Fig. 7C, and panels in Fig. 7D.

To further investigate how these two mutants handicapped SIE, we adopted a low-background bi-fluorescence complementation (BIFC) assay (27, 28) to examine the inter-molecular interactions among the mutant p28 proteins. This improved BIFC assay used a monomeric Venus (mVenus) protein that is split between the 10^th^ and 11^th^ (the last) β-strands to minimize self-association between the untagged fragments of this fluorescent protein, hence drastically reducing false positives. An additional advantage of this new system is that the large N-terminal mVenus fragment (NVen, 10 β-strands) could be readily detected using a GFP antibody. To prepare for the BIFC assay, the V181A and F182A mutants of p28, along with wildtype p28, were fused to the N-termini of NVen and CVen (the 11^th^ β-strand of mVenus). The resulting binary constructs were delivered into *N. benthamiana* cells to detect potential intermolecular interactions. As a negative BIFC control, we paired the NVen-tagged p28 with CVen-tagged p60 – a deletion mutant of p88 with its N-terminal p28 portion removed. p28 and p60 did not interact with each other in preliminary BIFC assays (also see later).

To ensure the constructs expressed the intended proteins, we first examined the accumulation levels of the NVen-tagged proteins (the CVen tagged forms could not be detected due to the small size of the tag). As shown in Fig. 7C, the wildtype p28-NVen accumulated to very high levels in the presence of p28-CVen, but to a much more modest level in the presence of p60-CVen (lanes 2 and 3), suggesting that it was stabilized by p28-p28 interactions (see later). Consistent with the results of Figs. 7A and B, substantially reduced accumulation was observed with both V181A-NVen and F182A-NVen proteins, and the reduction was more dramatic for F182A-NVen (Fig. 7C, lanes 4 and 5). Compared to the p28-NVen/p60-CVen negative control (lane 3), the p28-NVen protein was slightly increased by V181A-CVen but modestly reduced by F182A-CVen (lanes 6 and 8). Notably, presence of p28-CVen did not enhance the accumulation of either V181A-NVen or F182A-NVen (lanes 7 and 9), suggesting that their levels were regulated independently of the presence of wildtype p28.

When viewed under a confocal microscope, the p28-NVen/p28-CVen pair gave rise to intensely bright yellow foci that resembled the p28-GFP foci (Fig. 7D, panel 2). By contrast, the p28-NVen/p60-CVen pair was completely devoid of any yellow fluorescent signals, indicating an absence of p28-p60 interaction (panel 3). Interestingly, V181A-NVen/V181A-CVen still interacted to form foci in easily detectable, but visibly fewer numbers (panel 4). By contrast, the F182A-NVen/F182A-CVen co-delivery resulted in substantially fewer foci that were mostly smaller and dimmer (panel 5). However, pairing F182A-CVen with p28-NVen, or vice versa, partially restored the formation of bright yellow foci (panels 8 and 9). This experiment was repeated multiple times with similar results. The number of foci in each of the interactions was then quantified by counting the foci in multiple equal sized leaf sections containing 40-50 cells (see Materials and Methods for details), and plotting the counts on box plots (Fig. 7E). The quantification revealed that the V181A mutant proteins interacted with each other (Group #4), or wildtype p28 (Groups #6 and 7), to yield modestly fewer fluorescent foci than p28/p28 interactions (approximately 20% reduction). By contrast, the F182A/F182A interaction led to a reduction of approximately 80% in focus numbers (Group #5). Notably, this reduction was partially reversed by changing one of the interacting partners to a wildtype p28 derivative (Groups #8 and 9). Thus, the F182A mutant proteins interacted with each other very inefficiently, probably due to low protein concentrations, but were able to coalesce with the nucleated core formed by wildtype p28 to produce the large foci detectable with confocal microscopy. Importantly, the lower protein levels of V181A and F182A mutant p28 proteins, coupled with reduced numbers of p28 foci, strongly correlated with the weakened SIE elicitation by these mutant proteins.

## DISCUSSION

SIE protects host cells from being sequentially infected by the same virus. Although use of SIE in controlling plant virus diseases has been extensively documented, its underlying molecular mechanism, and evolutionary rationale, remain poorly understood (14). Using the model virus TCV, we have previously established that SIE occurred inside individual super-invaded cells, and acted to repress the replication of the superinfecting viral genomes (16, 29). We further discovered that SIE in TCV infections is mediated by a single TCV-encoded protein, the p28 auxiliary replication protein, likely through the formation of a self-perpetuating polymeric state that sequesters homologous p28 monomers. This model raised the possibility that SIE exists to ensure that progeny genomes of a virus are excluded from additional rounds of replication, thus minimizes error accumulation within single cells (18). By extension, cells invaded by many copies of the same virus simultaneously, frequently in the form of collective infectious units (30–33), likely also experience similar molecular events that ensure only a small number of the entered genomes is allowed to replicate, and be subjected to cellular level selections. If this is true, then viral mutants that weaken SIE are expected to relax this restriction, allowing more viral genomes to replicate for multiple cycles in the same cells, causing fast error accumulation in viral populations, and probably swift collapse of these populations.

The current study was designed to test the predictions of our working hypothesis by first identifying mutations within TCV p28 that compromise SIE. Our systematic deletion mutagenesis of p28, using two different C-terminally tagged forms of p28 as the backbones, allowed us to narrow down the SIE-eliciting activity of p28 to three regions: aa 2-50, 126-155, and 156-190. The aa 156-190 region was then further interrogated because deletion of this region caused a consistent loss of SIE in both p28-GFP and p28-HA backgrounds. These efforts ultimately led to the identification of two amino acid residues, namely valine (V) and phenylalanine (F) at positions 181 and 182, respectively, whose replacement by alanine (A) in replicons led to the decoupling of p28 replication function and its SIE elicitation function. It is interesting to note that these two residues appear to have different levels of conservation, with F182, but not V181, being highly conserved among the p28 analogs of multiple carmoviruses (not shown).

The lower protein levels of p28(V181A) correlated with weak SIE in both transient expressions and TCV replicons, suggesting that a protein concentration threshold had to be reached for p28 to elicit a robust SIE. This is consistent with the model proposed by us in earlier studies (16, 18). Why did the F182A mutant proteins, being even less abundant than V181A proteins, interfere with SIE elicitation only in the transiently expressed form, but not in replicons? The answer might lie in the inability of the F182A-containing replicon to replicate by itself. Specifically, replication of the F182A mutant replicon must rely on wildtype (or V181A) p28 encoded by a different replicon (Fig. 6), and occurred in a small number of cells (approximately 5%). In these cells, wildtype p28 protein was translated from the primary transcripts of the replicon construct, but the replication of the wildtype replicon was delayed (16). This delay created a precious window for the F182A mutant replicon to initiate replication with borrowed wildtype p28. Once succeeding in replicating itself, the F182A mutant p28 would be produced in massive amounts, thus neutralizing the lower accumulation levels. More importantly, the presence of wildtype p28 in these cells would facilitate the multimerization of p28 to form nucleation cores onto which the F182A p28 could coalesce, forming the large p28 foci that exert strong SIE. Indeed our BIFC experiments illustrated that the low self-interaction efficiency of the p28(F182A) protein was drastically improved when one of the interacting partners was changed to wildtype p28 (Fig. 7D). This scenario does not apply to the V181A mutation because the V181A replicon is capable of replicating itself. Replication of the V181A replicon could occur in some cells before the wildtype p28 translated from a co-introduced wildtype replicon reaches the SIE-eliciting concentration threshold. Conversely, wildtype TCV could still replicate in a fraction of these cells because the V181A mutant p28 protein could only exert SIE inefficiently.

To summarize, the current study identifies two single aa mutations in TCV p28 that differentially impaired the two opposite functions of p28 – replication and SIE elicitation, hence decoupling these two functions of p28. These mutations will allow us to investigate the impact of SIE in the rate of mutation accumulation in TCV populations, providing insights regarding the evolutionary rationale for preserving SIE functionality in virus infections. These investigations would in turn unravel potential targets for antiviral therapy, and prompt novel management strategies for virus diseases.

## MATERIALS AND METHODS

### Constructs

Constructs 2X35S::TCV_sg2R, p28-HA, p28 (tag-free), and Core35S::p88 (tag-free) were described in previous studies (16, 23). The 2X35S::p28-GFP construct was a slightly modified version of the previous p28-GFP. The new version replaced the NotI site (GCGGCCGCA) linking p28 and GFP coding sequences with a BamHI site plus three additional nts (GGATCCGGA). Nearly all deletion mutants, as well as single aa substitution mutants, were generated using synthetic gBlocks fragments (IDT, Coralville, IA) that were integrated into various backbone plasmids (e.g. p28-GFP, p28-HA, or TCV_sg2R) using Gibson Assembly cloning (NEB, Ipswich, MA). The identities of all new constructs were verified with Sanger sequencing. The constructs for the BIFC assay were assembled by fusing the NVen and CVen fusion tags to the C-termini of p28, p28(V181A), and p28(F182A), using as the 2X35S::p28-GFP construct backbone. The cDNA sequence of NVen and CVen was derived from the paper of Gookin and Assman (28).

### Agro-infiltration

Upon verification, all of the constructs were introduced into *Agrobacterium tumefaciens* strain C58C1 with electroporation (34, 35). To carry out the experiments described in the Results section, various combinations of *Agrobacterium* suspensions were mixed together and delivered into *N. benthamiana* leaves as described (23, 29, 34, 36). A p19-expressing *Agrobacterium* strain was included in all combinations to alleviate RNA silencing-mediated mRNA degradation.

### RNA extraction and Northern blotting

Total RNA was extracted from agro-infiltrated *N. benthamiana* leaves using the Direct-zol RNA Miniprep kit (Zymo Research, Irvine, CA). To ensure consistency, four equivalent leaf sections derived from infiltrated leaves of four different plants were pooled before RNA extraction. The RNA extraction procedure included a DNase treatment step that removes DNA contamination. The RNA was then quantified with NanoDrop and subjected to Northern blotting as described (23, 29, 34, 36).

### Protein extraction and Western blotting

Proteins were extracted from the agro-infiltrated N. benthamiana leaves following the protocol of Zhang et al (22). The polyclonal GFP and the monoclonal HA antibodies were purchased from Invitrogen. The AP-conjugated anti-rabbit (GFP) and anti-mouse (HA) secondary antibodies were purchased from Pierce and Novus, respectively. The HRP-conjugated anti-rabbit (GFP) secondary antibody was purchased from Abcam.

### Confocal microscopy and quantification of fluorescent cells

Confocal microscopic observations were carried out using a Leica Confocal microscope (TCS SP5) available at Molecular and Cellular Imaging Center at the Ohio Agricultural Research and Development Center, The Ohio State University. To quantify the percentage of cells expressing GFP, mCherry, or reconstituted mVenus (Fig. 5 and 7), the agro-infiltrated *N. benthamiana* leaf sections were photographed under the confocal microscope, at the low magnification (10X), and for three separate channels (GFP/mVenus, mCherry, and merged). For each treatment, five independent leaf sections were photographed, resulting in 15 images. These images, typically encompassing 40-50 cells, were then divided into 100 equal subdivisions with the grid line feature of the PhotoShop software. Subdivisions that are more than 50% filled with a given fluorescence (GFP, mCherry, or both) were counted as positive for the respective protein, the total counts of which represent a percentage in a given image. For each treatment five counts were obtained, and standard deviation were computed using these counts. In the case of BIFC experiments, the number of cells in each of the selected leaf section was estimated using a co-expressed ER-mCherry protein (not shown). The reconstituted mVenus foci was counted for each of the five leaf sections, and the counts were quantified using box plots.

## ACKNOWLEDGMENTS

We thank the labs of Drs. Lucy Stewart and Peg Redinbaugh for generous equipment sharing, and other members of the Qu lab for stimulating discussions. This study was supported by a grant from the National Science Foundation (#1758912), a SEEDS grant from the Ohio Agricultural Research and Development Center, Graduate Assistantships from OSU and OARDC to Q.G. and R.S., respectively, as well as tuition assistances to S.Z., R.S., and Q.G. from the Department of Plant Pathology, OSU.

## LITERATURE

1. Nethe M, Berkhout B, van der Kuyl AC. 2005. Retroviral superinfection resistance. Retrovirology 2:52.

2. Schaller T, Appel N, Koutsoudakis G, Kallis S, Lohmann V, Pietschmann T, Bartenschlager R. 2007. Analysis of Hepatitis C Virus Superinfection Exclusion by Using Novel Fluorochrome Gene-Tagged Viral Genomes. J Virol 81:4591.

3. Tscherne DM, Evans MJ, von Hahn T, Jones CT, Stamataki Z, McKeating JA, Lindenbach BD, Rice CM. 2007. Superinfection exclusion in cells infected with hepatitis C virus. J Virol 81:3693–3703.

4. Zou G, Zhang B, Lim P-Y, Yuan Z, Bernard KA, Shi P-Y. 2009. Exclusion of West Nile Virus Superinfection through RNA Replication. J Virol 83:11765.

5. Simon KO, Cardamone JJ, Whitaker-Dowling PA, Youngner JS, Widnell CC. 1990. Cellular mechanisms in the superinfection exclusion of vesicular stomatitis virus. Virology 177:375–379.

6. Julve JM, Gandía A, Fernández-del-Carmen A, Sarrion-Perdigones A, Castelijns B, Granell A, Orzaez D. 2013. A coat-independent superinfection exclusion rapidly imposed in Nicotiana benthamiana cells by tobacco mosaic virus is not prevented by depletion of the movement protein. Plant Mol Biol 81:553–564.

7. Dietrich C, Maiss E. 2003. Fluorescent labelling reveals spatial separation of potyvirus populations in mixed infected Nicotiana benthamiana plants. J Gen Virol 84:2871–2876.

8. Bergua M, Zwart MP, El-Mohtar C, Shilts T, Elena SF, Folimonova SY. 2014. A viral protein mediates superinfection exclusion at the whole-organism level but is not required for exclusion at the cellular level. J Virol 88:11327–11338.

9. Miyashita S, Kishino H. 2010. Estimation of the size of genetic bottlenecks in cell-to-cell movement of soil-borne wheat mosaic virus and the possible role of the bottlenecks in speeding up selection of variations in trans-acting genes or elements. J Virol 84:1828–1837.

10. Takahashi T, Sugawara T, Yamatsuta T, Isogai M, Natsuaki T, Yoshikawa N. 2007. Analysis of the Spatial Distribution of Identical and Two Distinct Virus Populations Differently Labeled with Cyan and Yellow Fluorescent Proteins in Coinfected Plants. Phytopathology 97:1200–1206.

11. Folimonova SY. 2012. Superinfection Exclusion Is an Active Virus-Controlled Function That Requires a Specific Viral Protein. J Virol 86:5554.

12. Zhou X, Sun K, Zhou X, Jackson AO, Li Z. 2019. The matrix protein of a plant rhabdovirus mediates superinfection exclusion by inhibiting viral transcription. J Virol JVI.00680-19.

13. Ziebell H, Carr JP. 2010. Cross-Protection, p. 211–264. In Advances in Virus Research. Elsevier.

14. Zhang X-F, Qu F. 2016. Cross Protection of Plant Viruses: Recent Developments and Mechanistic Implications, p. 241–250. In Wang, A, Zhou, X (eds.), Current Research Topics in Plant Virology. Springer International Publishing, Cham.

15. Folimonova SY. 2013. Developing an understanding of cross-protection by Citrus tristeza virus. Front Microbiol 4:76.

16. Zhang X-F, Sun R, Guo Q, Zhang S, Meulia T, Halfmann R, Li D, Qu F. 2017. A self-perpetuating repressive state of a viral replication protein blocks superinfection by the same virus. PLOS Pathog 13:e1006253.

17. Tatineni S, French R. 2016. The Coat Protein and NIa Protease of Two Potyviridae Family Members Independently Confer Superinfection Exclusion. J Virol 90:10886–10905.

18. Zhang X-F, Zhang S, Guo Q, Sun R, Wei T, Qu F. 2018. A New Mechanistic Model for Viral Cross Protection and Superinfection Exclusion. Front Plant Sci 9.

19. White KA, Skuzeski JM, Li W, Wei N, Morris TJ. 1995. Immunodetection, Expression Strategy and Complementation of Turnip Crinkle Virus p28 and p88 Replication Components. Virology 211:525–534.

20. Cao M, Ye X, Willie K, Lin J, Zhang X, Redinbaugh MG, Simon AE, Morris TJ, Qu F. 2010. The Capsid Protein of Turnip Crinkle Virus Overcomes Two Separate Defense Barriers To Facilitate Systemic Movement of the Virus in Arabidopsis. J Virol 84:7793–7802.

21. Zhang X, Zhang X, Singh J, Li D, Qu F. 2012. Temperature-Dependent Survival of Turnip Crinkle Virus-Infected Arabidopsis Plants Relies on an RNA Silencing-Based Defense That Requires DCL2, AGO2, and HEN1. J Virol 86:6847–6854.

22. Qu F, Ye X, Morris TJ. 2008. Arabidopsis DRB4, AGO1, AGO7, and RDR6 participate in a DCL4-initiated antiviral RNA silencing pathway negatively regulated by DCL1. Proc Natl Acad Sci 105:14732–14737.

23. Zhang S, Sun R, Guo Q, Zhang X-F, Qu F. 2019. Repression of turnip crinkle virus replication by its replication protein p88. Virology 526:165–172.

24. Shaner NC, Steinbach PA, Tsien RY. 2005. A guide to choosing fluorescent proteins. Nat Methods 2:905–909.

25. Powers JG, Sit TL, Qu F, Morris TJ, Kim K-H, Lommel SA. 2008. A Versatile Assay for the Identification of RNA Silencing Suppressors Based on Complementation of Viral Movement. Mol Plant Microbe Interact 21:879–890.

26. May J, Johnson P, Saleem H, Simon AE. 2017. A Sequence-Independent, Unstructured Internal Ribosome Entry Site Is Responsible for Internal Expression of the Coat Protein of Turnip Crinkle Virus. J Virol 91:e02421–16.

27. Kodama Y, Hu C-D. 2010. An improved bimolecular fluorescence complementation assay with a high signal-to-noise ratio. BioTechniques 49:793–805.

28. Gookin TE, Assmann SM. 2014. Significant reduction of BiFC non-specific assembly facilitates in planta assessment of heterotrimeric G-protein interactors. Plant J 80:553–567.

29. Zhang X-F, Guo J, Zhang X, Meulia T, Paul P, Madden LV, Li D, Qu F. 2015. Random Plant Viral Variants Attain Temporal Advantages During Systemic Infections and in Turn Resist other Variants of the Same Virus. Sci Rep 5.

30. Chen Y-H, Du W, Hagemeijer MC, Takvorian PM, Pau C, Cali A, Brantner CA, Stempinski ES, Connelly PS, Ma H-C, Jiang P, Wimmer E, Altan-Bonnet G, Altan-Bonnet N. 2015. Phosphatidylserine Vesicles Enable Efficient En Bloc Transmission of Enteroviruses. Cell 160:619–630.

31. Mirabelli C, Wobus CE. 2018. All Aboard! Enteric Viruses Travel Together. Cell Host Microbe 24:183–185.

32. Andreu-Moreno I, Sanjuán R. 2018. Collective Infection of Cells by Viral Aggregates Promotes Early Viral Proliferation and Reveals a Cellular-Level Allee Effect. Curr Biol 28:3212-3219.e4.

33. Miyashita S, Ishibashi K, Kishino H, Ishikawa M. 2015. Viruses Roll the Dice: The Stochastic Behavior of Viral Genome Molecules Accelerates Viral Adaptation at the Cell and Tissue Levels. PLOS Biol 13:e1002094.

34. Qu F, Ren T, Morris TJ. 2003. The Coat Protein of Turnip Crinkle Virus Suppresses Posttranscriptional Gene Silencing at an Early Initiation Step. J Virol 77:511–522.

35. Qu F, Ye X, Hou G, Sato S, Clemente TE, Morris TJ. 2005. RDR6 Has a Broad-Spectrum but Temperature-Dependent Antiviral Defense Role in Nicotiana benthamiana. J Virol 79:15209–15217.

36. Zhang X-F, Sun R, Guo Q, Zhang S, Meulia T, Halfmann R, Li D, Qu F. 2017. A self-perpetuating repressive state of a viral replication protein blocks superinfection by the same virus. PLOS Pathog 13:e1006253.

